# BldD-based bimolecular fluorescence complementation for *in vivo* detection of the second messenger cyclic di-GMP

**DOI:** 10.1101/2021.08.26.457767

**Authors:** Manuel Halte, Mirka E. Wörmann, Maxim Bogisch, Marc Erhardt, Natalia Tschowri

**Author notes:** These authors contributed equally to this work (Manuel Halte, Mirka E. Wörmann). Competing interest statement: The authors declare no competing interests.

## Abstract

The widespread bacterial second messenger bis-(3’-5’)-cyclic diguanosine monophosphate (c-di-GMP) is an important regulator of biofilm formation, virulence and cell differentiation. C-di-GMP-specific biosensors that allow detection and visualization of c-di-GMP levels in living cells are key to our understanding of how c-di-GMP fluctuations drive cellular responses. Here, we describe a novel c-di-GMP biosensor, CensYBL, that is based on c-di-GMP-induced dimerization of the effector protein BldD from *Streptomyces* resulting in bimolecular fluorescence complementation of split-YPet fusion proteins. As a proof-of-principle, we demonstrate that CensYBL is functional in detecting fluctuations in intracellular c-di-GMP levels in the Gram-negative model bacteria *Escherichia coli* and *Salmonella enterica* serovar Typhimurium. Using deletion mutants of c-di-GMP diguanylate cyclases and phosphodiesterases, we show that c-di-GMP dependent dimerization of CBldD-YPet results in fluorescence complementation reflecting intracellular c-di-GMP levels. Overall, we demonstrate that the CensYBL biosensor is a user-friendly and versatile tool that allows to investigate c-di-GMP variations using single-cell and population-wide experimental set-ups.

**Importance:** The second messenger c-di-GMP controls various bacterial functions including development of resistant biofilm communities and transition into dormant spores. *In vivo* detection of c-di-GMP levels is therefore crucial for a better understanding of how intracellular c-di-GMP levels induce changes of bacterial physiology. Here, we describe the design of a novel c-di-GMP biosensor and demonstrate its effective application in investigating fluctuations in intracellular c-di-GMP levels in *Escherichia coli* and *Salmonella enterica* serovar Typhimurium on a population-based and single-cell level.

## Introduction

Cyclic dinucleotide second messengers are key components of signal transduction systems for bacterial adaptation to environmental cues. Bis-(3′,5′)-cyclic di-guanosine-monophosphate (c-di-GMP) was discovered in 1987 as a stimulator of cellulose synthase in *Komagataeibacter xylinus* (Ross *et al*., 1987) and is now recognized as a conserved central regulator of bacterial physiology (Jenal *et al*., 2017). Diguanylate cyclases (DGCs) and c-di-GMP-specific phosphodiesterases (PDEs) possess antagonistic activities and control intracellular c-di-GMP levels in response to diverse signals. DGCs synthesize c-di-GMP out of GTP through their catalytic GGDEF domains that are named after key amino acids in their active sites (Paul *et al*., 2004). c-di-GMP-specific PDEs carry either EAL or HD-GYP domains. EAL-type PDEs hydrolyze c-di-GMP to the linear 5′-phosphoguanylyl-(3′-5′)-guanosine (pGpG) dinucleotide (Christen *et al*., 2005), while HD-GYP domains cleave c-di-GMP to two molecules of GMP (Bellini *et al*., 2014).

c-di-GMP exerts a global regulatory role on diverse bacterial functions, including biofilm formation, motility, virulence, cell cycle progression and cell differentiation by binding to an array of protein effectors and riboswitches (Chou and Galperin, 2016; Römling *et al*., 2013). Intracellular levels of the molecule range from nanomolar to low-micromolar concentrations between bacterial species (Abel *et al*., 2013; Sarenko *et al*., 2017). Interestingly, c-di-GMP levels can also vary between individual cells of an isogenic population leading to phenotypic diversity, as exemplified by *Salmonella* Typhimurium during infection (Petersen *et al*., 2019). In *Bacillus subtilis* subpopulations, high c-di-GMP levels correlate with the transition to sporulation, while low levels are associated with competence (Weiss *et al*., 2019). Moreover, in *Caulobacter crescentus* asymmetrical distribution of c-di-GMP within the cell through local action of c-di-GMP-metabolizing enzymes was reported to be critical for extracellular organelle positioning (Christen *et al*., 2010).

Monitoring c-di-GMP patterns in living cells revealed fundamental new insights into c-di-GMP-dependent phenotypic heterogeneity and *in vivo* functions of DGCs and PDEs. This became only possible by the application of fluorescence-based methods for detection of c-di-GMP *in vivo*. Three different approaches can be employed for biosensing of the second messenger. First, c-di-GMP-responsive promoters can be fused to fluorescent proteins and used as reporters of the dinucleotide in living cells. A transcriptional fusion of the promoter of the *cdrA* gene, encoding a large adhesin in *Pseudomonas aeruginosa*, to the gene encoding for green fluorescent protein (GFP) revealed that *P. aeruginosa* responds to surface attachment by increased production of c-di-GMP (Rodesney *et al*., 2017; Rybtke *et al*., 2012). Second, riboswitches that specifically bind c-di-GMP to their aptamer domains (Sudarsan *et al*., 2008) can be utilized as c-di-GMP biosensors. Ligand-binding aptamers were successfully fused to RNA aptamers that incorporate small fluorophores like 3,5-difluoro-4-hydroxybenzylidene imidazolinone (DFHBI) and emit fluorescence. In such riboswitch-based biosensors, binding of c-di-GMP leads to conformational changes in the fused aptamer which results in DFHBI binding and fluorescence that can be monitored to estimate c-di-GMP levels (Kellenberger *et al*., 2013; Nakayama *et al*., 2012). Alternatively, c-di-GMP-responsive riboswitches can be fused to fluorescent proteins to detect the second messenger in living cells (Weiss *et al*., 2019; Zhou *et al*., 2016). Finally, c-di-GMP-binding proteins can be utilized as biosensors for the dinucleotide. For example, the c-di-GMP effector YcgR from *Salmonella enterica* serovar Typhimurium undergoes a conformational change upon ligand binding which can be coupled to Förster resonance energy transfer (FRET) (Christen *et al*., 2010; Kulasekara *et al*., 2013).

In this study, we describe the development and application of a c-di-GMP biosensor that utilizes bimolecular fluorescence complementation (BiFC) of the yellow fluorescence protein (YPet) upon ligand binding to the c-di-GMP effector BldD. The principle is based on two split non-fluorescent fragments of YPet, which, when brought together by interaction between proteins or proteins domains that are fused to each fragment, undergo fluorescence complementation (Hu *et al*., 2002). The transcriptional regulator BldD has recently been identified as a c-di-GMP effector in *Streptomyces venezuelae* (Tschowri *et al*., 2014). During their developmental life cycle, *Streptomyces* undergo a complex physiological and morphological transition that culminates in the formation of dormant spores (Bush *et al*., 2015). BldD binds a tetrameric c-di-GMP to the RxD-X8-RxxD signature motif in the C-terminal domain of BldD (CBldD), which in turn drives BldD dimerization. Dimerized BldD binds to its target promoters to represses a broad regulon of sporulation genes (den Hengst *et al*., 2010). Structural and biochemical studies revealed that the BldD_2_-(c-di-GMP)_4_ complex assembles in an ordered, sequential manner. First, a c-di-GMP dimer binds to BldD motif 2 (RxxD), which induces conformational changes that favor binding of the second dimer to motif 1 (RxD) (Schumacher *et al*., 2017). Importantly, in the BldD_2_-(c-di-GMP)_4_ complex, the two CBldDs are separated by about 10 Å and are linked together purely by c-di-GMP that acts as a macromolecular dimerizer. Therefore, c-di-GMP-dependent dimerization of CBldD is ideally suited to be employed for c-di-GMP driven BiFC and thus, for *in vivo* detection of c-di-GMP. Here, we describe the design and demonstrate successful application of a biosensor based on bimolecular fluorescence complementation of a YPet fusion to the c-di-GMP effector BldD (CensYBL) to quantify and monitor fluctuations in c-di-GMP levels of individual cells and bacterial populations of the model organisms *Escherichia coli* and S. Typhimurium.

## Results

### Design and characterization of CensYBL as *in vivo* c-di-GMP biosensor

Guided by structural and biochemical data revealing that CBldD dimerization is driven purely by c-di-GMP, we sought to design a CBldD-based *in vivo* sensor for c-di-GMP that utilizes the bimolecular fluorescence complementation principle. The BiFC approach is based on complementation of split fluorophores and is frequently used for the detection of protein-protein interactions (Kerppola, 2009). First, we split the yellow fluorescence protein into two fragments and tested protein-protein interaction-driven fluorescence complementation by using an established two-hybrid interaction (Karimova *et al*., 1998). For that, the N-terminal fraction of YPet (NYPet), comprising amino acids 1-173, was fused to the dimerizing leucine zipper part (“zip”) domain of the yeast transcription factor GCN4 in the pKT25 vector and the C-terminal part of YPet (CYPet), comprising amino acids 156-239, was fused to the zip domain in pUT18C. As a control, we designed pKT25 and pUT18C-based vectors expressing either NYPet or CYPet without being fused to the interacting Zip proteins (Figure S1 A). As shown in Figure S1 B, expression of NYPet-Zip1 and CYPet-Zip2 in *E. coli* led to a fluorescence YPet signal, while no fluorescence was detectable in cells co-expressing the separated YPet fragments only.

Next, we fused NYPet and CYPet, respectively, to the C-terminal domain of BldD from *S. venezuelae* (CBldD, comprising residues 80-166) including a GSGGG linker between the two domains and cloned the two synthetically generated open reading frames in tandem into the pTrc99a-FFA vector (Amann *et al*., 1988) (Figure 1 A-C). We named the biosensor fusion protein CensYBL for <u>C</u>-di-gmp s<u>ENS</u>or <u>Y</u>pet <u>BL</u>dD. For normalization of CensYBL expression between individual cells, we generated a transcriptional fusion of the *mcherry* gene downstream of the CensYBL-encoding genes. As such, in the plasmid pCensYBL, expression of the three genes *nypet::cbldD, cypet::cbldD* and *mcherry* is controlled by the same IPTG inducible P*trc* promoter. In line with our expectations, we detected strong YPet and mCherry signals, respectively, upon induction of pCensYBL for 1 h using 250 µM IPTG in *E. coli* (Figure 2 A).

**Figure 1:**
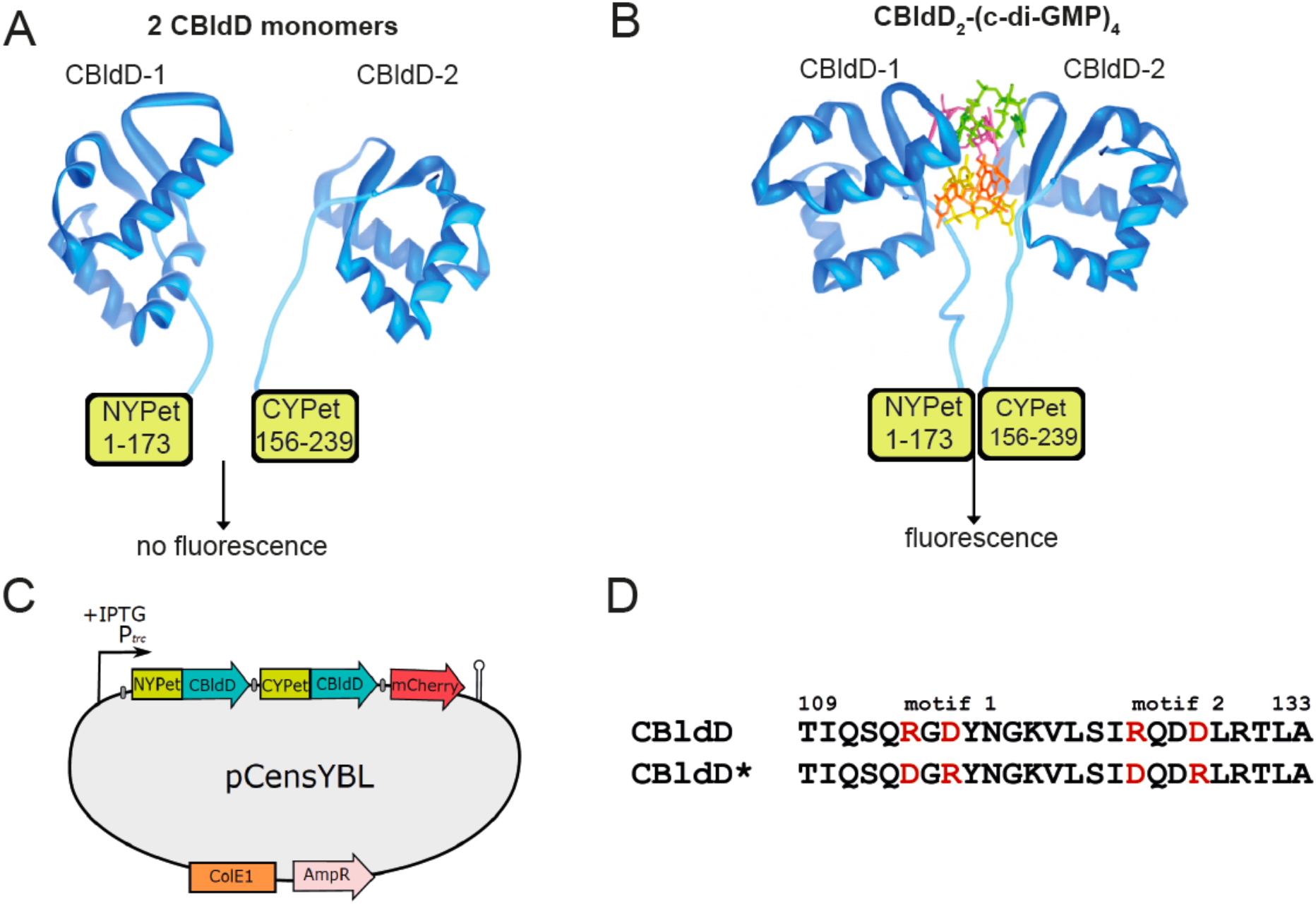
Design principle of the CensYBL c-di-GMP biosensor. (A-B) Schematic illustration of the CensYBL c-di-GMP biosensor fusion proteins that consist of either the N-terminal part of YPet (NYPet; amino acids 1-173) or the C-terminal part of YPet (CYPet; amino acids 156-239) connected to the C-terminal c-di-GMP-binding domain of BldD (CBldD). (A) In absence of c-di-GMP, CBldD adopts a monomeric conformation that does not support bimolecular complementation of the split YPet fluorophore. (B) Upon binding of a tetrameric c-di-GMP, BldD dimerizes, which allows complementation of the two YPet fragments leading to a fluorescence signal. (C) The pTrc99a-FFA-based pCensYBL vector harbors a *nypet::cbldD, cypet::cbldD* and *mcherry* operon under the control of the IPTG inducible P*trc* promoter. D. Protein alignment of a BldD fragment (amino acids 109 - 133) comprising the c-di-GMP binding motif 1 (RxD) and motif 2 (RxxD) in CBldD that are essential for c-di-GMP binding. The amino acids highlighted in red were mutagenized to DxR and DxxR in CBldD* to abolish c-di-GMP binding.

**Figure 2:**
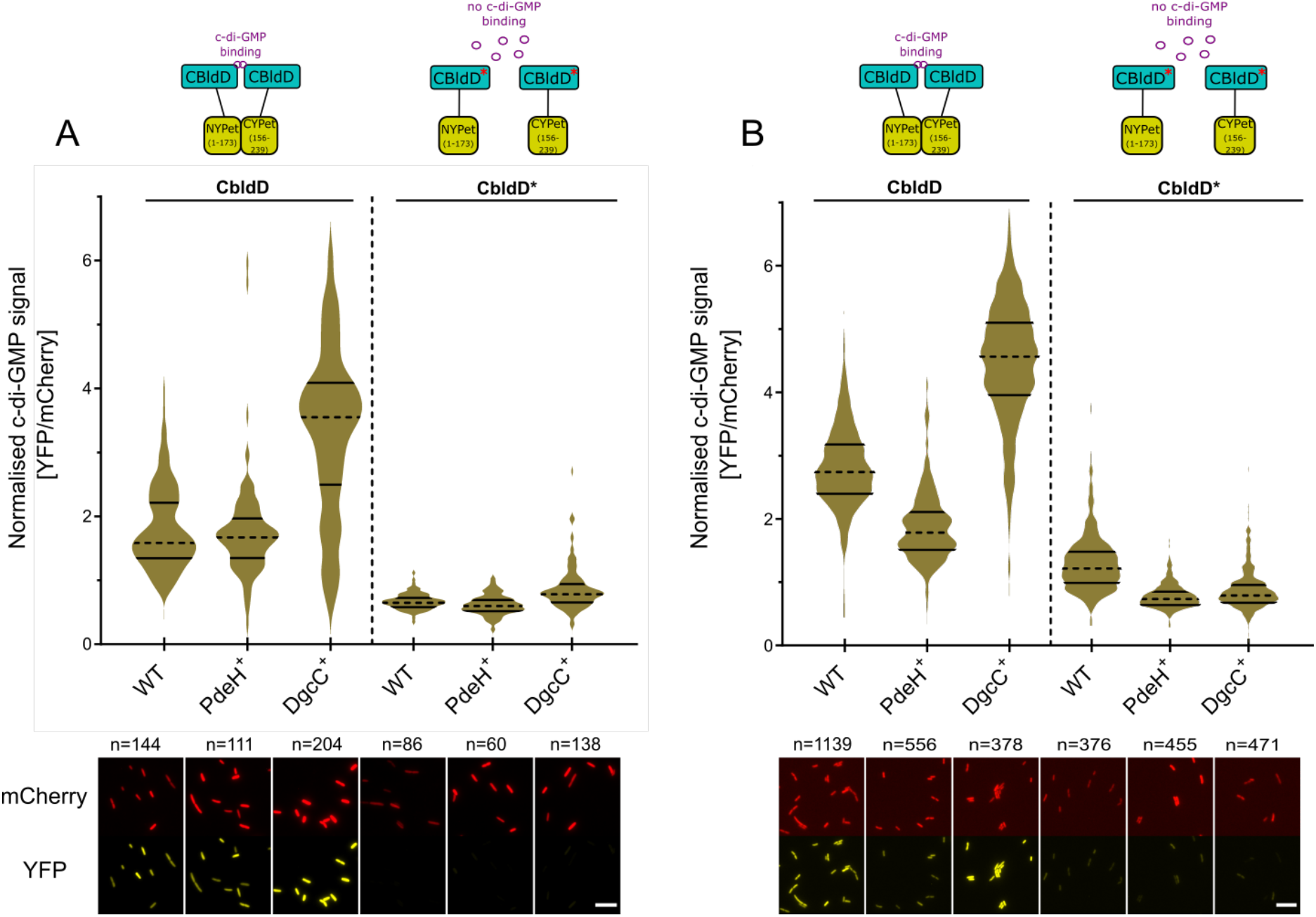
The CensYBL biosensor detects changes of intracellular c-di-GMP levels. The active version of the CensYBL biosensor (CbldD) or the inactive version (CbldD*) was used to quantify intracellular c-di-GMP levels *E. coli* (A) and *S*. Typhimurium (B). Strains overexpressing the PDE PdeH or the DGC DgcC were investigated using fluorescence microscopy. The top part of the panels shows violin plots of the normalized c-di-GMP signal obtained from single cell measurements. The dashed lines represent the median and solid lines the quartiles. Representative fluorescent microscopy images of the mCherry and YFP channels are shown in the lower part of the panels. Scale bar = 10µm.

CBldD interacts with c-di-GMP using two adjacent motifs. Motif 1 is composed of residues 114-116 (RxD) and motif 2 comprises residues 125-128 (RxxD) (Figure 1 D). Mutagenesis of either motif 1 or motif 2 completely abolishes c-di-GMP-binding (Tschowri *et al*., 2014). We aimed to design an inactive BldD-based biosensor that is blind for c-di-GMP as a specificity control (CensYBL*). For that, we swapped the arginine residues in motif 1 and 2 with aspartic acid and *vice versa* as described in Tschowri et al., 2014 leading to CbldD*, which is unable to bind the second messenger. As shown in Figure 2 A, expression of the split YPet fragments fused to CbldD* led to weak background fluorescence only. Altogether, our data demonstrate that c-di-GMP dependent CBldD-dimerization results in functional complementation of NYPet and CYPet and can be utilized as a tool for *in vivo* detection of c-di-GMP.

Upon confirming successful complementation of split YPet fused to CbldD when expressed from pCensYBL, we assessed the *in vivo* protein expression levels of CYPet- and NYPet-CbldD and mCherry in *E. coli*-K12 W3110 (Hayashi *et al*., 2006). Active and inactive form of CensYBL were expressed in strains with modulated levels of c-di-GMP due to overexpression of a PDE (PdeH^+^) or a DGC (DgcC^+^) from the IPTG-inducible pCAB18 plasmid (Tschowri *et al*., 2009). We next performed Western Blot analyses of whole cell lysates using anti-BldD and anti-mCherry antibodies, respectively (Figure S2 A). In line with our expectations, no striking differences of NYPet/CYPet-CbldD and mCherry protein levels were observed in the wild type strain compared to the strains expressing a PDE/DGC, confirming that differences in c-di-GMP levels do not impact the expression of CensYBL and mCherry. Respective loading controls are shown in Figure S2 B. However, we detected lower protein levels for the CensYBL* inactive biosensor. Split YFP were reported to be prone to aggregation when interaction between the split fragments failed (Kostecki *et al*., 2010). This is also the case for CensYBL* fusions and likely causes protein aggregation or instability. Accordingly, we concluded that CensYBL expression is independent of the intracellular c-di-GMP levels, and that the observed changes in YPet intensities depend on the intracellular c-di-GMP concentration.

### CensYBL-based detection of c-di-GMP in individual *E. coli* and *S*. Typhimurium cells

We then aimed to evaluate CensYBL *in vivo* to monitor intracellular c-di-GMP levels. For this, we used *E. coli* and *S*. Typhimurium strains carrying plasmids overexpressing the PDE PdeH or the DGC DgcC (Pesavento *et al*., 2008; Richter *et al*., 2020). Both enzymes are well-characterized and affect c-di-GMP levels in an antagonistic manner. PdeH is a stand-alone EAL domain phosphodiesterase previously known as YhjH in *E. coli* and STM3611 in *S*. Typhimurium. DgcC, formerly designated as YaiC in *E. coli* and STM0385 / AdrA in *S*. Typhimurium, is a GGDEF domain DGC. In both species, those enzymes are involved in biofilm regulation by a complex regulatory network controlled by c-di-GMP (Crepin *et al*., 2017; Kader *et al*., 2006; Pesavento *et al*., 2008; Richter *et al*., 2020). By using those well-characterized enzymes, we aimed to modify the intracellular c-di-GMP levels in order to assess CensYBL sensitivity and its capability to measure different c-di-GMP concentrations on a single-cell level.

Strains harboring plasmids overexpressing PdeH or DgcC under the control of a P*lac* promoter, in combination with plasmids encoding the active or inactive CensYBL biosensor, were grown to exponential phase under IPTG inducing conditions. We then determined the YFP and mCherry fluorescence of individual cells using live-cell epifluorescence microscopy. The mCherry fluorescence signal was used for normalization of expression, and fluorescence levels of YFP were calculated relative to the mCherry levels for each cell. We found that in the *E. coli* DgcC^+^ strain, the median c-di-GMP signal (YFP/mCherry fluorescence ratio) was increased by 2-fold (Figure 2 A). Contrary to our expectations, the c-di-GMP signal observed in the PdeH^+^ strain was similar to the wild type. We note, however, that a c-di-GMP concentration of ~60 nM was reported for *E. coli* wild type cells (Sarenko *et al*., 2017). As the K_D_ of BldD-CTD for c-di-GMP was determined to be about 2.5 µM (Tschowri *et al*., 2014), it is possible that we are not able to detect minor differences in intracellular c-di-GMP levels between the wild type and upon PdeH^+^ overexpression in *E. coli*. We note that for the CensYBL* mutant biosensor defective in c-di-GMP binding, we observed a background fluorescence signal only in all strains analyzed. In complementing experiments, we next tested the ability of the CensYBL biosensor to monitor intracellular c-di-GMP levels in *S*. Typhimurium strains carrying the same plasmids overexpressing a PdeH or DgcC, together with the active/inactive CensYBL biosensor (Figure 2 B). We detected a similar increase of the intracellular c-di-GMP signal in a strain expressing DgcC^+^ compared to the respective *E. coli* strain. Contrary to our data obtained with *E. coli*, we observed a 1.5-fold decrease in the c-di-GMP signal in the PdeH^+^ strain compared to wild type. We conclude that CensYBL can detect intracellular differences in c-di-GMP concentrations in Gram-negative bacteria and can be used for functional analysis of DGCs and PDEs *in vivo*.

### CensYBL-based detection of c-di-GMP fluctuations in a *S. enterica* population

In addition to quantifying intracellular c-di-GMP levels using microscopy-based single cell analysis, we aimed to determine if the CensYBL biosensor can be used for c-di-GMP detection and quantification using plate reader measurements. Such an approach would be advantageous to facilitate the screening of a large number of mutants in a time-efficient and easy manner. We therefore measured the c-di-GMP signal in *Salmonella* strains overexpressing either DgcC or PdeH as presented above, as well as using mutants deleted either for the PDE-encoding genes *pdeH* or *pdeC* and the Δ*pdeH* Δ*pdeC* double mutant. PdeC (STM4264) is a PDE phosphodiesterase ortholog to *E. coli* YjcC and contains an EAL domain involved in invasion and biofilm formation control in *Salmonella* (Ahmad *et al*., 2017; Ahmad *et al*., 2011).

After induction of CensYBL expression, the cells were collected by centrifugation, washed in PBS and loaded onto 96-well plates. Measured fluorescence values of YFP and mCherry were divided by the OD600, and subsequent YFP/OD600 values were made relative to mCherry/OD600 levels (see Material and Methods). We observed a 0.9-fold decrease and a 3.3-fold increase in the c-di-GMP signals of the PdeH^+^ and DgcC^+^ strains (Figure 3 A), respectively, similar as observed using the microscopy-based approach. For the single deletion mutants of *pdeH* or *pdeC*, we observed a 1.21 and 1.10-fold increase in the c-di-GMP signals, respectively, while the double deletion mutant (Δ*pdeH* Δ*pdeC*) displayed a 1.5-fold signal increase compared to the wild type (Figure 3 B). The inactive CensYBL* biosensor displayed a low signal compared to the active biosensor that was similar between the strains tested. Altogether, our data demonstrate that CensYBL biosensor can be utilized for *in vivo* detection and quantification of the intracellular c-di-GMP level of individual bacteria and for screening of c-di-GMP mutants using plate reader measurements.

**Figure 3:**
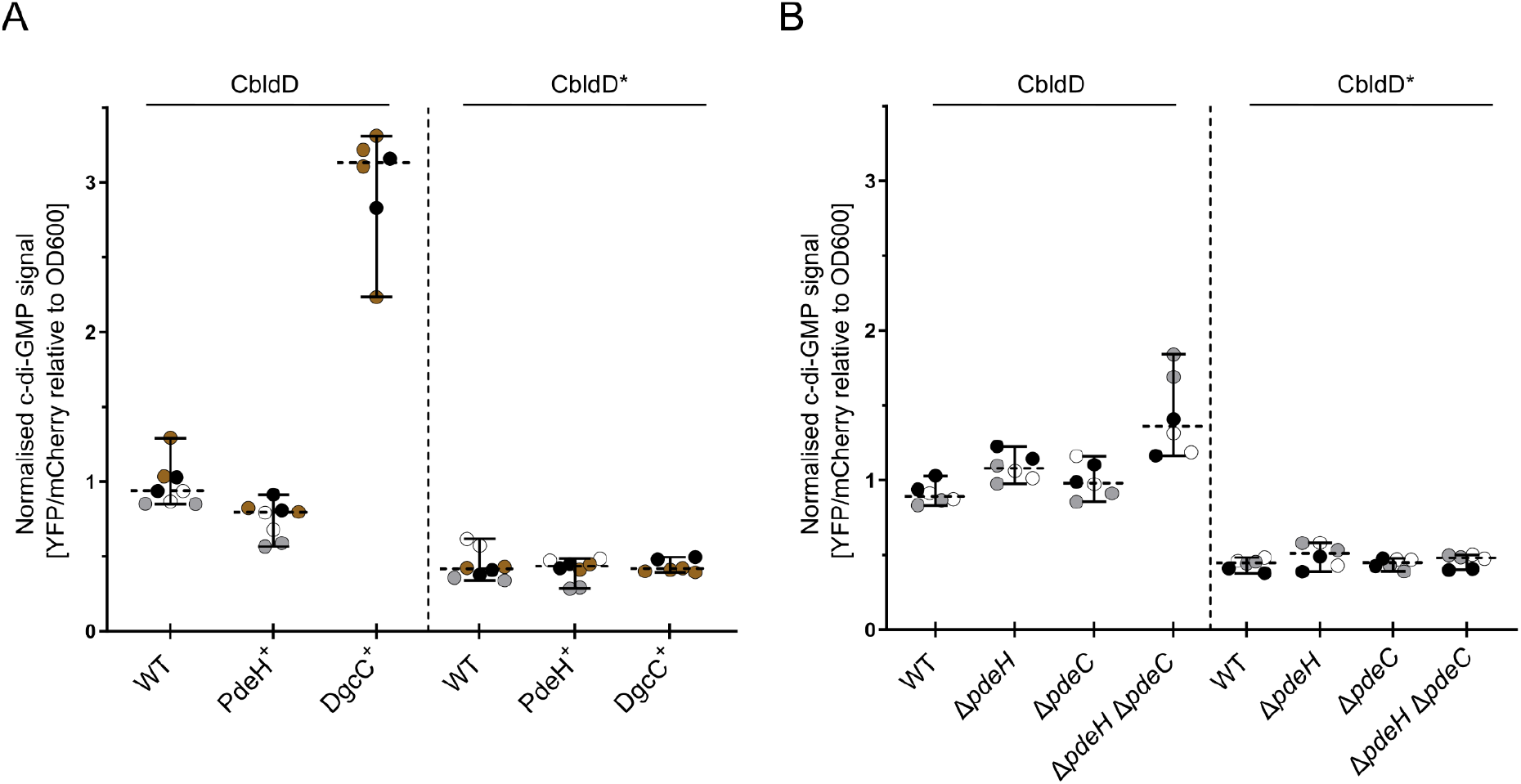
The CensYBL c-di-GMP biosensor detects fluctuations in c-di-GMP levels in population measurements. The active version of the CensYBL biosensor (CbldD) or the inactive version (CbldD*) was used to quantify c-di-GMP levels *E. coli* (A) and *S*. Typhimurium (B) using plate reader-based population measurements. (A) C-di-GMP levels were quantified in *E. coli* strains overexpressing the PDE PdeH or the DGC DgcC. The dashed lines represent the median of the normalized c-di-GMP signals and the solid lines the 95% CI, obtained from six biological replicates (the mean of each biological replicate was calculated from three technical replicates measurements). (B) C-di-GMP levels were quantified in *S*. Typhimurium PDE deletion strains (Δ*pdeH* / Δ*pdeC*). The dashed lines represent the median of the normalized c-di-GMP signals and the solid lines the 95% CI, obtained from six biological replicates analyzed on three different days (the mean of each biological replicate was calculated from three technical replicates measurements). The color represents the biological replicates of each day.

## Discussion

In this work, we developed a fluorescent c-di-GMP biosensor, CensYBL, and demonstrate that it can detect fluctuations in intracellular c-di-GMP levels in the Gram-negative species *E. coli* and *S*. Typhimurium. The CensYBL biosensor exploits the mode-of-action of the c-di-GMP binding domain of BldD, a transcriptional regulator that orchestrates the transition from hyphae to spore in *Streptomyces* (Tschowri *et al*., 2014). BldD forms a homo-dimeric complex, where two BldD monomers dimerize in presence of four c-di-GMP molecules. Hereby, the second messenger specifically interacts with the RxD-X8-RxxD motif in the C-terminal domain of BldD and glues together the otherwise separated BldD monomers. We reasoned that this dimerization mechanism, that is driven exclusively by c-di-GMP, is perfectly suited for utilization as a c-di-GMP sensor system. For the design of such a BldD-based biosensor, we therefore fused the c-di-GMP-sensing domain of BldD to the C-terminal and N-terminal regions of YPet in order to utilize BiFC as readout for c-di-GMP binding. Our data shown in Figures 2 and 3 demonstrate that the CensYBL biosensor is suitable to detect changes in the intracellular c-di-GMP concentration both using fluorescence microscopy at the single cell level, and using plate reader-based population measurements, which highlights the potential of CensYBL for versatile applications studying c-di-GMP regulation.

As a proof-of-principle, we investigated the ability of the CensYBL biosensor to detect differences in intracellular c-di-GMP levels upon overexpression of PDEs and DGCs. Interestingly, we found that the CensYBL biosensor did not detect a lower c-di-GMP signal upon overproduction of PdeH in *E. coli* when compared to the wild type strain (Figure 2 A). The physiological c-di-GMP levels in *E. coli* were previously estimated to be in the ~60 nM range (Sarenko *et al*., 2017) and are likely even lower upon overexpression of PdeH. Since the c-di-GMP binding affinity of BldD has been reported to be ~2.5 µM (Tschowri *et al*., 2014), we cannot exclude that the low c-di-GMP concentrations in wild type *E. coli* may represent the lower detection limit of the CensYBL biosensor. We note, however, that overproduction of DgcC resulted in a strong increase of the c-di-GMP signal detected by CensYBL (Figure 2 A).

A non-active version of our CensYBL, expressing a mutated CBldD* in which the c-di-GMP-binding signature was modified to DxR-X8-DxxR, allowed us to demonstrate that bimolecular fluorescence complementation of our biosensor was specifically dependent on c-di-GMP binding to CBldD. Notably, we observed decreased BldD protein levels in the non-active version of CensYBL* (Figure S2 A), which we attributed to the lack of CBldD*-NYPet and CBldD*-CYPet interaction likely leading to protein instability. A similar observation was made by Kostecki et al, while studying the protein-protein interaction of *E. coli* DmsA with its binding partner DmsD, using YPet bimolecular fluorescence complementation (Kostecki *et al*., 2010).

We further note that the mechanism if and how the BldD_2_-(c-di-GMP)_4_ complex dissociates is unknown and therefore the CensBYL biosensor might currently not be ideally suited to monitor transient and dynamic changes of c-di-GMP levels. Future versions of CensYBL might include a C-terminal degradation tag as a target for tail-specific proteases to prevent accumulation of stable, complemented CensYBL biosensor, similar to the use of unstable GFP variants for investigations of transient changes in gene expression (Andersen *et al*., 1998).

Over the years, various fluorescence and bioluminescence-based sensors have been developed to detect c-di-GMP levels. Translational fusions of fluorophores to YcgR, a c-di-GMP receptor, were used to detect c-di-GMP binding using FRET microscopy (Christen *et al*., 2010), a technique that requires a specific expertise. Moreover, FRET microscopy might result in lower fluorescence signals and unwanted cross-talk between the donor and acceptor fluorophore when compared to conventional fluorescent microscopy of a single fluorescent protein. Further, transcriptional fusions of fluorescent proteins to c-di-GMP regulated promoters were used to obtain insights into the mechanism of c-di-GMP regulation of *P. aeruginosa* after surface attachment (Rodesney *et al*., 2017; Rybtke *et al*., 2012). However, these transcriptional fusions are limited to *Pseudomonas* and strictly c-di-GMP dependent promoters might not be known for a given model organism. Finally, riboswitches that specifically bind c-di-GMP were utilized to detect c-di-GMP levels by fusing RNA aptamers to those structures. Binding of c-di-GMP to the riboswitch results in a conformational change and fluorescence emission after addition of small fluorophores ligands (Kellenberger *et al*., 2013; Nakayama *et al*., 2012). However, those systems require a high concentration of fluorescent ligands and washing steps before imaging, which may interfere with physiological processes and might not be suitable for time-resolved investigations of dynamic changes in c-di-GMP levels.

Altogether, the CensYBL c-di-GMP biosensor reported in this study provides the c-di-GMP field with a user-friendly and sensitive system to measure and quantify intracellular c-di-GMP. We expect that this new tool will greatly facilitate future investigations of the complex regulation and physiological importance of the c-di-GMP second messenger in bacteria.

## Material and Methods

### Bacterial strains, plasmids and oligonucleotides

Strains, plasmids and oligonucleotides used in this study are listed in Table S1. The construction of the plasmids pKT25_*nypet::cbldD*/*cbldD\** and pUT18C_*cypet::cbldD*/*cbldD\** is described in the supplementary experimental procedures. The active and inactive versions of CensYBL (pTrc99A-FFA_*ypet*(split)-*cbldD[cbldD*]*_*mCherry*_*rrnB*) were generated using Gibson Assembly of two fragments. The pTrc99A-FAA backbone sequence was amplified by inverse PCR using primers MW182/183. The sequence coding for the active and inactive biosensor, respectively, was amplified from plasmids pSS170_Perm*_*ypet*(split)-*cbldD /cbldD\**_*mCherry*_*rrnB* (for details see supplementary experimental procedures) using primers MW180/181. pSS09 and pSS88 served as templates for amplification of the coding region of *ypet* and *mcherry*, respectively. The sequence of the active and inactive biosensor can be found in the supplementary data S1.

### Fluorescence microscopy

Strains harboring the active and inactive version of the biosensor were grown in LB at 37 °C or 30 °C, respectively, for 45 min before induction with 250 µM IPTG. Cultures growth was resumed for 1h and 2h, respectively for *E. coli* and *S*. Typhimurium, cells were harvested, washed in PBS, and applied on 1% agarose pads (Sigma). Slides were imaged using a Zeiss Axio Observer Z1 inverted epifluorescence microscope. Images were acquired at 63× magnification with an Axiocam 506 mono CCD-camera using the Zen 2.6 pro software using the filter sets 46 He (YFP) and 64 He (mCherry) and exposure time of 600 ms (YFP) and 800 ms (mCherry), respectively.

### Image analysis

Fluorescent microscopy images were processed using Fiji (Schindelin *et al*., 2012). Fluorescence levels were quantified using Fiji plugin MicrobeJ (Ducret *et al*., 2016). After segmentation and detection of the cells using phase contrast images, the signal-to-noise ratio of mCherry and YFP was measured directly by MicrobeJ in each cell after detection. The subsequent YFP/mCherry ratio was then manually calculated for each cell. Values obtained were plotted using GraphPad Prism version 9.

### Plate reader measurements

Cultures were grown and induced as described above for the fluorescence microscopy experiments. After growth, 2 mL cells were collected by centrifugation at 15.000×g and the cell pellet was resuspended in 500 µL PBS. Each well of a black 96-well, clear bottom plate was loaded with 150 µL of resuspended culture (Corning, Product number 3603). Measurements of OD (600 nm) and fluorescence were performed using a TECAN Infinite M200 plate reader (TECAN) using excitation/emission wavelengths of 590/620 nm and 510/540 nm for mCherry and YFP, respectively. For each sample, the measured YFP and mCherry fluorescence signal was normalized to the OD600 signal. The YFP values were then normalized to the mCherry values to report the c-di-GMP signal for each sample and plotted using GraphPad Prism version 9.

## Acknowledgements

This work was supported in part by the European Research Council (ERC) under the European Union’s Horizon 2020 research and innovation program (grant to Marc Erhardt; agreement no. 864971). Research in Natalia Tschowri’s lab is funded by the DFG Emmy Noether-Program (TS 325/1-1) and the DFG Priority Program SPP 1879 (TS 325/2-2). We thank Heidi Landmesser and Kristin Funke for expert technical assistance.

## Author contributions

N.T. and M.E. designed the study. All authors designed and interpreted experiments, which were performed by M.H., M.E.W., and M.B. The figures were made by N.T. and M.H. The paper was written by M.H., M.E. and N.T. with input from the other authors.

