## Supplemental Data for "BldD-based bimolecular fluorescence complementation for *in vivo* detection of the second messenger cyclic di-GMP"

#### Supplementary experimental procedures

##### Plasmid construction

Plasmids pKT25_*nypet_zip*, pUT18C_*cypet_zip*, pKT25_*nypet* and pUT18C_*cypet* were constructed to analyze fluorescence complementation of split YPet fragments. Plasmid pKT25_*nypet_zip* was obtained by amplifying the 5´sequence of *ypet,* coding for the first 173 amino acids, from plasmid pSS09 using primer pair MW1/4. The *zip* sequence was amplified from plasmid pKT25_zip using primer pair MW73/74 and the *rrnB* terminator sequence was amplified from plasmid pMMB_HS_Bc3-5 using primer pair MW75/59. The resultant products were fused by splice overlap extension PCR (SOE PCR) (Horton *et al.*, 1989) with primer pair MW1/59. The pKT25 plasmid backbone was amplified by inverse PCR from plasmid pKT25_*zip* using primer pair MW2/56. Plasmid backbone and fused PCR fragments were assembled using Gibson assembly (Gibson *et al.*, 2009). The resultant plasmid was transformed into XL1BLue yielding strain XL1Bue_pKT25_*Nypet_zip* (NT147). Plasmid pKT25_*nypet* was constructed by inverse PCR from pKT25_*nypet_cbldD* using primer pair MW77/78. The PCR fragment was religated and introduced into XL1BLue yielding strain XL1Bue_pKT25_*nypet* (NT148). Plasmid pUT18C_*cypet_zip* was obtained by amplifying the 3´sequence of *ypet,* coding amino acids 156-239, from plasmid pSS09 using primer pair MW5/81. The *zip* sequence was amplified from plasmid pUT18C_*zip* using primer pair MW73/79 and the *rrnB* terminator sequence was amplified from plasmid pMMB_HS_Bc3-5 using primer pair MW80/59. The resultant products were fused SOE PCR with primer pair MW5/59. The pUT18C plasmid backbone was amplified by inverse PCR from plasmid pUT18C_*zip* using primer pair MW6/61. Plasmid backbone and fused PCR fragments were assembled using Gibson assembly (Gibson *et al.*, 2009). The resultant plasmids were transformed into XL1BLue yielding strain XL1Bue_pUT18C_*cypet_zip* (NT149). Plasmid pUT18C_*cypet* was constructed by inverse PCR from pUT18C_*cypet_cbldD* using primer pair MW82/78. The PCR fragment was religated and introduced into XL1BLue yielding strain XL1Blue_pUT18C_*cypet* (NT150). Plasmids pKT25_*nypet_zip* and pUT18C_*cypet_zip* and plasmids pKT25_*nypet* and pUT18C_*cypet* were cotransformed into W3110 resulting in strains NT169 and NT170, respectively.

Plasmids pKT25_*nypet::cbldD* and pKT25_*nypet::cbldD** were obtained by amplifying the *ypet* coding region comprising amino acids 1 - 173 from plasmid pSS09 using primer pair MW1/58. The relevant *bldD* sequence*,* encoding the C-terminal domain of the protein, was amplified from plasmid pET15b_*bldD* or pET15b_*bldD** (Tschowri *et al.*, 2014)using primer pair MW57/52. The *rrnB* terminator sequence was amplified from plasmid pMMB_HS_Bc3-5 with primer pair MW53/54. The resultant products were fused by splice overlap extension PCR using primers MW1/54. The pKT plasmid backbone was amplified by inverse PCR from plasmid pKT25_zip using primer pair MW2/56. Plasmid backbone and fused PCR fragments were assembled using Gibson assembly. The resultant plasmids were transformed into XL1BLue yielding strains XL1Bue pKT25_*nypet::cbldD* (strain collection number NT137) and XL1Blue pKT25_*nypet::cbldD** (strain collection number NT138) for storage.

Plasmids pUT18C_*cypet::cbldD* and pUT18C_*cypet::cbldD** were constructed by amplifying the *ypet* coding region covering amino acids 156 - 239 from plasmid pSS09 using primer pair MW5/63. Primer pair MW62/52 was used to amplify the 3′ region of *bldD* encoding CBldD from pET15b_*bldD* or pET15b_*bldD**, respectively. Using the primer pair MW53/59, the *rrnB* terminator sequence was amplified from plasmid pMMB_HS_Bc3-5. The resultant products were fused by SOE PCR using primers MW5/59. The pUT18C plasmid backbone was amplified by inverse PCR from plasmid pUT18C_zip using primer pair MW6/61. Plasmid backbone and fused PCR fragments were assembled using Gibson assembly and the resultant plasmids were transformed into XL1Blue yielding strains XL1Blue pUT18C_*cypet::cbldD* (strain collection number NT141) and XL1Blue pUT18C_*cypet::cbldD** (strain collection number NT142) respectively.

Plasmids pSS170_Perm*_*ypet*(split)-*cbldD_mCherry_rrnB* and pSS170_Perm*_*ypet*(split*)*-*cbldD*_mCherry_rrnB* were constructed for future application in *Streptomyces venezuelae*.

The sequences coding for NYpet-CBldD fusions with active or inactive CBldD were amplified from plasmids pKT25_*nypet_cbldD* and pKT_*nypet_cbldD** using primer pair MW140/141. The sequences coding for CYpet-CBldD fusions containing active or inactive CBldD were amplified from pUT18C_*cypet_cbldD* and pUT18C_*Cypet_CbldD** using primer pair MW142/143. The *mcherry* sequence was amplified from plasmid pSS88 and the *rrnB* terminator sequence was amplified from pMM_HS_Bc3-5 with primer pairs MW144/128 and MW129/136 respectively. The pSS170 plasmid backbone and the Perm* promoter sequence were amplified by inverse PCR from plasmid pSS170_Perm*_*nypet_cbldD* with primer pair MW135/139 and the five PCR fragments were assembled by Gibson assembly. The resultant plasmids were electroporated into DH5α resulting into strains DH5α_ pSS170_Perm*_*ypet*(split)-*CbldD_mCherry_rrnB* (NT238) and DH5α_ pSS170_Perm*_*ypet*(split)-*CbldD*_mCherry_rrnB* (NT239).

**Supplementary information**

**Table S1. Strains, plasmids and oligonucleotides used in this study**

| | **Genotype or comments** | **Source or reference** |
|---|---|---|
| **Strains** | | |
| W3110 | K-12 derivative; *F-, λ-, rpoS(Am), rph-1, Inv(rrnD-rrnE)* | (Hayashi *et al.*, 2006) |
| TH437 | *Salmonella enterica* serovar Typhimurium LT2 | J. Roth / K.T. Hughes |
| EM11244 | Δ*pdeC::FCF* (STM4264) | This study |
| EM11313 | Δ*pdeC::FCF* (STM4264) Δ*pdeH::FKF* | This study |
| EM11367 | Δ*pdeH::FKF* | This study |
| EM10758 | LT2 / pEM10758 (pTrc99A-FF4-Nypet-CbldD-Cypet-CbldD-mCherry, AmpR) | This study |
| EM10759 | LT2 / pEM10759 (pTrc99A-FF4-Nypet-CbldD*-Cypet-CbldD*-mCherry, [inactive version] AmpR) | This study |
| EM10860 | pEM10758 (pTrc99A-FF4-Nypet-CbldD-Cypet-CbldD-mCherry, AmpR) / pEM10858 (pCAB18-*pdeH*, CmR) | This study |

| | | |
|---|---|---|
| EM10861 | pEM10759 (pTrc99A-FF4-Nypet-CbldD*-Cypet-CbldD*-mCherry,AmpR) / pEM10858 (pCAB18-*pdeH*, CmR) | This study |
| EM10862 | pEM10758 (pTrc99A-FF4-Nypet-CbldD-Cypet-CbldD-mCherry, AmpR) / pEM10859 (pCAB18-*dgcC*, CmR) | This study |
| EM10863 | pEM10759 (pTrc99A-FF4-Nypet-CbldD*-Cypet-CbldD*-mCherry, AmpR) / pEM10859 (pCAB18-*dgcC*, CmR) | This study |
| EM11385 | Δ*pdeH::FKF* / pEM10758 (pTrc99A-FF4-Nypet-CbldD-Cypet-CbldD-mCherry, AmpR) | This study |
| EM11386 | Δ*pdeH::FKF* / pEM10759 (pTrc99A-FF4-Nypet-CbldD*-Cypet-CbldD*-mCherry, [inactive version] AmpR) | This study |
| EM11387 | Δ*pdeC::FCF* (STM4264) / pEM10758 (pTrc99A-FF4-Nypet-CbldD-Cypet-CbldD-mCherry, AmpR) | This study |
| EM11388 | Δ*pdeC::FCF* (STM4264) / pEM10759 (pTrc99A-FF4-Nypet-CbldD*-Cypet-CbldD*-mCherry, [inactive version] AmpR) | This study |
| EM11389 | Δ*pdeC::FCF* (STM4264) Δ*pdeH::FKF* / pEM10758 (pTrc99A-FF4-Nypet-CbldD-Cypet-CbldD-mCherry, AmpR) | This study |
| EM11390 | Δ*pdeC::FCF* (STM4264) Δ*pdeH::FKF* / pEM10759 (pTrc99A-FF4-Nypet-CbldD*-Cypet-CbldD*-mCherry, [inactive version] $Amp^R$) | This study |
| NT650 | W3110 pTrc99A-FFA_*ypet*(split)-*cbldD_mCherry_rrnB* (pCensYBL active); $Amp^R$ | This study |
| NT651 | W3110 pTrc99A-FFA_*ypet*(split)-*cbldD*_mCherry_rrnB* (pCensYBL inactive); $Amp^R$ | This study |
| NT652 | W3110 pTrc99A-FFA_*ypet*(split)-*cbldD_mCherry_rrnB* (pCensYBL active) + pCAB18::*pdeH*; $Amp^R$ , $Cm^R$ | This study |

| | | |
|---|---|---|
| NT653 | W3110 pTrc99A-FFA_*ypet*(split)-*cbldD_mCherry_rrnB* (pCensYBL active) + pCAB18::*dgcC*; $Amp^R$ , $Cm^R$ | This study |
| NT654 | W3110 pTrc99A-FFA_*ypet*(split)-*cbldD*_mCherry_rrnB* (pCensYBL inactive) + pCAB18::*pdeH*; $Amp^R$ , $Cm^R$ | This study |
| NT655 | W3110 pTrc99A-FFA_*ypet*(split)-*cbldD*_mCherry_rrnB* (pCensYBL inactive) + pCAB18::*dgcC*; $Amp^R$ , $Cm^R$ | This study |
| NT169 | W3110 pKT25_*nypet::zip* + pUT18C_*cypet::zip*, $Kan^R$ , $Amp^R$ | This study |
| NT170 | W3110 pKT25_*nypet* + pUT18C_*cypet*, $Kan^R$ , $Amp^R$ | This study |

| **Plasmids** | | |
|---|---|---|
| pKT25 | Low copy vector encoding the T25 fragment of *Bordetella pertussis cyaA* downstream of the MCS; $Kan^R$ | Euromedex |
| pUT18C | High copy vector encoding the T18 fragment of *B. pertussis cyaA* downstream of the MCS; $Amp^R$ | Euromedex |
| pSS09 | Codon-optimized *ypet* in pMS82 | unpublished |
| pSS88 | Codon-optimized *mcherry* in pIJ10257 | (Schlimpert *et al.*, 2017) |
| pMMB_HS_Bc3-5 | source for *rrnB* terminator sequence | (Zamorano-Sanchez *et al.*, 2019) |
| pKD3 | frt-*cat*-frt vector plasmid, $Cm^R$ | (Datsenko and Wanner, 2000) |
| pKD46 | Temperature sensitive plasmid for LambdaRed recombination, $Amp^R$ | (Datsenko and Wanner, 2000) |
| NT137 | pKT25_*nypet::cbldD*, $Kan^R$ | This study |

| | | |
|---|---|---|
| NT138 | pKT25_*nypet::cbldD**, Kan$^R$ | This study |
| NT141 | pUT18C_*cypet::cbldD*, Amp$^R$ | This study |
| NT142 | pUT18C_*cypet::cbldD**, Amp$^R$ | This study |
| NT147 | pKT25_*nypet::zip*, Kan$^R$ | This study |
| NT149 | pUT18C_*cypet::zip*, Amp$^R$ | This study |
| NT148 | pKT25_*nypet*, Kan$^R$ | This study |
| NT150 | pUT18C_*cypet*, Amp$^R$ | This study |
| NT291 / pCensYBL (active) | pTrc99A-FFA_*ypet*(split)-*cbldD_mCherry_rrnB* | This study |
| NT292 pCensYBL (inactive) | pTrc99A-FFA_*ypet*(split)-*CbldD*_mCherry_rrnB* | This study |
| NT428 | pCAB18::*dgcC*; Cm$^R$ | (Tschowri *et al.*, 2009) |
| NT429 | pCAB18::*pdeH*; Cm$^R$ | (Tschowri *et al.*, 2009) |

| Oligonucleotide | Sequence |
|---|---|
| **Oligonucleotides used for cloning of pKT25_*nypet::cbldD/cbldD**** | |
| MW1: F_pKT25_*ypet*(1-173) | AAACAGCTATGGTCTCCAAGGGCGAGGAGCTGTTCACC |
| MW58: R-linkGSGSS_*Nypet* | GCTCGCCGCCGCCGGAGCCCTCGATGTTGTGGCGGATCTTGAAG |
| MW57: F_*Nypet*_linkGSGGG_*CbldD* | TCGAGGGCTCCGGCGGCGGCGAGCCGCCGCCGAAGCTCGTCC |
| MW52: R_*rrnB*term_*CbldD* | GTGGGACCACCGTCAGTTCTCCTCGTGGGCGACG |
| MW53: | GAGGAGAACTGACGGTGGTCCCACCTGACCCC |

| | |
|---|---|
| F_*CbldD_rrnB*term | |
| MW54: R_pKT_*rrnB*term | CCGAATTCTTAGTCACGGGAAATCCGTGTGATTGTTGCCG |
| MW2: R_*ypet*(1-173)_pKT25 | GAGACCATAGCTGTTTCCTGTGTGAAATTGTTATCCGC |
| MW56: F_*rrnB*term_pKT | GATTTCCCGTGACTAAGAATTCGGCCGTCGTTTTACAAC |
| **Oligonucleotides used for cloning of pKT25_*nypet::zip*** | |
| MW1: F_pKT25_*ypet*(1-173) | AAACAGCTATGGTCTCCAAGGGCGAGGAGCTGTTCACC |
| MW4 R_*zip*_(linker)*ypet*(1-173): | GTACCCGGAACTCGCCGGAGCCGGCGGCGGAGCCGGCGGAGCCCTCGATGTTGTGGCGGATCTTGAAGTTGGC |
| MW73: F_*zip*_shortlink | TCCGGCGGCGGCCGGGTACCTATCCAGCGTATG |
| MW74: R_*zip*(pKT)_*rrnB*term | GTGGGACCACCGTTACTTAGGTACCCCACGTTCAC |
| MW75: F_*rrnB*term_*zip*(pKT) | GTACCTAAGTAACGGTGGTCCCACCTGACCC |
| MW59: R_pUT_*rrnB*term | CATATTACTTAGTCACGGGAAATCCGTGTGATTGTTGCCG |
| MW2: R_*ypet*(1-173)_pKT25 | GAGACCATAGCTGTTTCCTGTGTGAAATTGTTATCCGC |
| MW56: F_*rrnB*term_pKT | GATTTCCCGTGACTAAGAATTCGGCCGTCGTTTTACAAC |
| **Oligonucleotides used for cloning of pUT18C_*cypet::zip*** | |
| MW5: F_pUT18C_*ypet*(156-239) | AAACAGCTATGGACAAGCAGAAGAACGGCATCAAGGCC |
| MW81: | GATAGGTACCCGGCCGCCGCCGGAGCCCTTGTAC |

| | |
|---|---|
| R_*Cypet*_shortlink_*zip* | |
| MW73: F_*zip*_shortlink | TCCGGCGGCGGCCGGGTACCTATCCAGCGTATG |
| MW79: R_*zip*(pUT)_*rrnB*term | GTGGGACCACCGTTATATCGATGAATTCGAGCTCGGTAC |
| MW80: F_*rrnB*term_*zip*(pUT) | TCATCGATATAACGGTGGTCCCACCTGACCCCATG |
| MW59: R_pUT_*rrnB*term | CATATTACTTAGTCACGGGAAATCCGTGTGATTGTTGCCG |
| MW6: R_*ypet*(156-239)_pUC18 | TTGTCCATAGCTGTTTCCTGTGTGAAATTGTTATCCGC |
| MW61: F_*rrnB*term_pUT | GATTTCCCGTGACTAAGTAATATGGTGCACTCTCAGTACAATCTGC |
| **Oligonucleotides used for cloning of pKT25_*nypet*** | |
| MW77: R_inv.PCR_pKT_*Nypet* | GGACCACCGTTAGCCGCCGCCGGAGCCCTCGATG |
| MW78: F_inv.PCR_pKT_*Nypet* | TCCGGCGGCGGCTAACGGTGGTCCCACCTGACCCCATGC |
| **Oligonucleotides used for cloning of pUT18C_*cypet*** | |
| MW78: F_inv.PCR_pKT_*Nypet* | TCCGGCGGCGGCTAACGGTGGTCCCACCTGACCCCATGC |
| MW82: R_inv.PCR_pUT_Cypet | GGACCACCGTTAGCCGCCGCCGGAGCCCTTGTAC |
| **Oligonucleotides used for cloning of pUT18C_*cypet::cbldD/cbldD**** | |
| MW5: F_pUT18C_*ypet*(156-239) | AAACAGCTATGGACAAGCAGAAGAACGGCATCAAGGCC |
| MW63: R_linkGSGGG_*Cypet* | GCTCGCCGCCGCCGGAGCCCTTGTACAGCTCGTTCATGCCCTC |

| | |
|---|---|
| MW62: F_*Cypet*_linkGSGGG_ *CbldD* | ACAAGGGCTCCGGCGGCGGCGAGCCGCCGCCGAAGCTCGTCC |
| MW52: R_*rrnB*term_*CbldD* | GTGGGACCACCGTCAGTTCTCCTCGTGGGCGACG |
| MW53: F_*CbldD_rrnB*term | GAGGAGAACTGACGGTGGTCCCACCTGACCCC |
| MW59: R_pUT_*rrnB*term | CATATTACTTAGTCACGGGAAATCCGTGTGATTGTTGCCG |
| MW6: R_ypet(156-239)_pUC18 | TTGTCCATAGCTGTTTCCTGTGTGAAATTGTTATCCGC |
| MW61: F_rrnBterm_pUT | GATTTCCCGTGACTAAGTAATATGGTGCACTCTCAGTACAATCTGC |
| **Oligonucleotides used for cloning of pTrc99A-FFA_*ypet*(split)-*cbldD/cbldD**_*mCherry_rrnB* (pCensYBL)** | |
| MW182: F_inv.PCR_pTrc99a_FFA | CCGTGAAAGCTTGGCTGTTTTGGCGGATGAGAGAAGATTTTC |
| MW183: R_inv.PCR_pTrc99a_FFA | GGAGACCATATGCTGTTTCCTGTGTGAAATTGTTATCCGCTCAC |
| MW180: F_5ýpet_BS_GA | CACAGGAAACAGCATATGGTCTCCAAGGGCGAGGAGCTG |
| MW181: R_3´rrnB_BS_GA | GCCAAAACAGCCAAGCTTTCACGGGAAATCCGTGTGATTGTTGC |
| **Oligonucleotides used for cloning of pSS170_Perm*_*ypet*(split)-*cbldD/cbldD**_*mCherry_rrnB*** | |
| MW140: F_BS_Nypet_CbldD_GA | GCCCGTCATATGGTCTCCAAGGGCGAGGAGCTGTTC |
| MW141: R_BS_Nypet_CbldD_G | TCATATCCTCCTTCCGGACCGATTTATCAGTTCTCCTCGTGGGCGACG |

| | |
|---|---|
| A | |
| MW142: F_BS_Cypet_CbldD_GA | AATCGGTCCGGAAGGAGGATATGAATATGGACAAGCAGAAGAACGGCATCAAG |
| MW143: R_BS_Cypet_CbldD_GA | AAATGACCTCCTCGATAGTCGCATTATCAGTTCTCCTCGTGGGCGACG |
| MW144: F_BS_mCherry_GA | ATGCGACTATCGAGGAGGTCATTTTTATGGTCTCCAAGGGCGAGGAGG |
| MW128: R_BS_mCherry | GTGGGACCACCGTTATCACTTGTACAGCTCGTCCATGCCG |
| MW129: F_BS_rrnB | TACAAGTGATAACGGTGGTCCCACCTGACCCCATG |
| MW136: R_BS_rrnB_GA | TTTATCAAGCTTTCACGGGAAATCCGTGTGATTGTTGC |
| MW135: F_pSS170_BS_GA | GATTTCCCGTGAAAGCTTGATAAACTTATCATCCCCTTTTGCTGATGG |
| MW139: R_pSS170_BS_GA | GCCCTTGGAGACCATATGACGGGCCTCCTGTTCTAGACG |
| **Oligonucleotides used for *ΔpdeC::FCF* construction** | |
| 5'-STM4264-FCF-fw | cccatcgtttacttgagtataaatctgatattatcaaaaaGTGTAGGCTGGAGCTGCTTC |
| 3'-STM4264-FCF-rev | gcgtatcgtcagaggcgcgggcgtttgttgtagccaggcgCATATGAATATCCTCCTTAG |
| 5'-STM4264-check-fw | CTATATCACAAAAACCAGCT |
| 3'-STM4264-check-rev | ACATTAGGCGAGTATATTCG |

### Figure S1

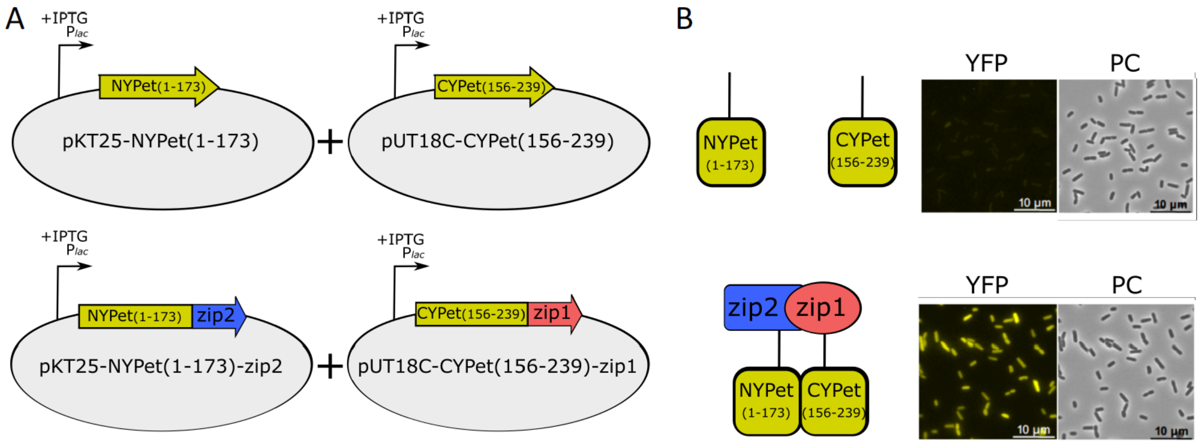


**Figure S1: Protein interaction dependent fluorescent complementation of split YPet fragments to a functional fluorophore.** (A) The N-terminal part of YPet (NYPet; amino acids 1-173) and the C-terminal part of YPet (CYPet; amino acids 156-239) were expressed in *E. coli*-K12 W3110 from pKT25 or pUT18C, respectively, either alone or fused to the leucine zipper region (Zip) of the yeast protein GCN4 (Karimova *et al.*, 1998). (B) The interaction of the Zip protein fusions resulted in fluorescence complementation of YPet (microscopy images lower right side).

**Figure S2**

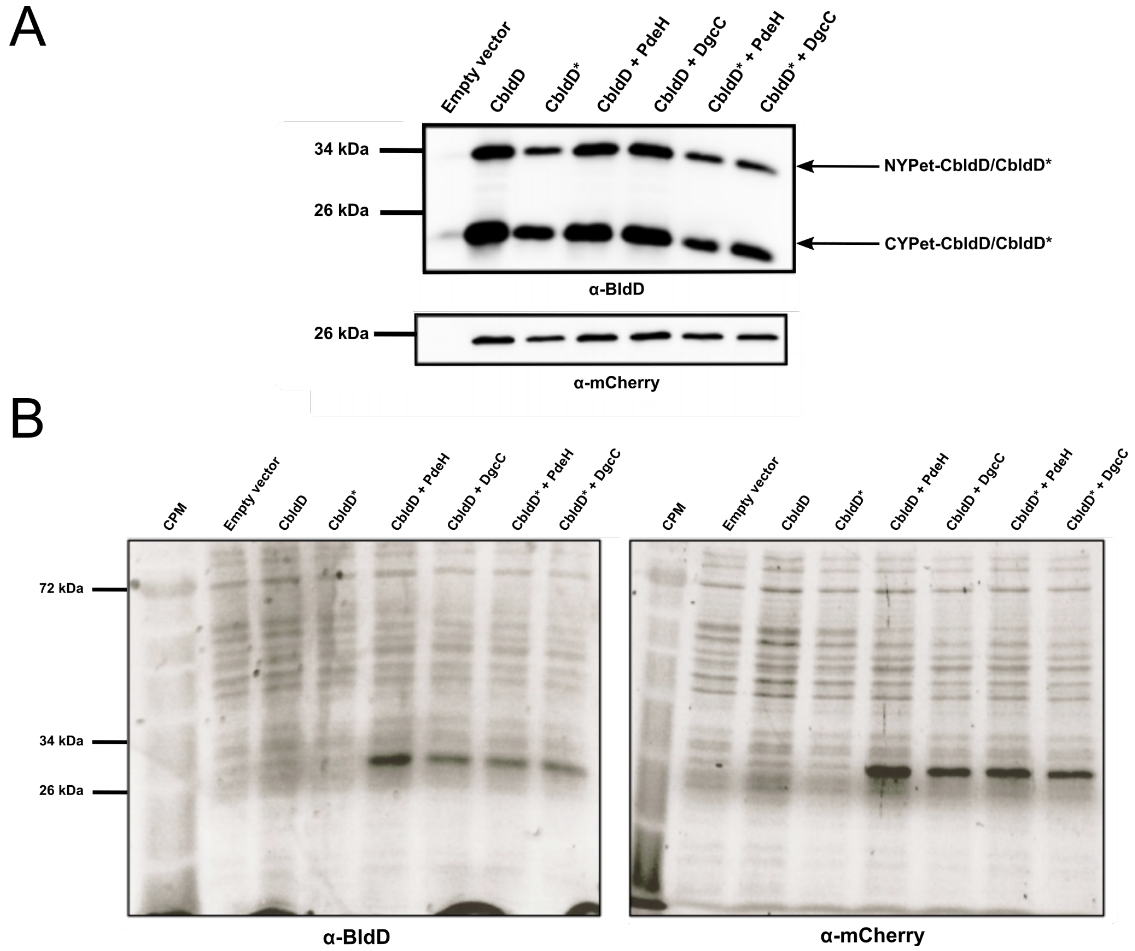


**Figure S2: Western blot analysis of CensYBL biosensor fusion proteins in *E. coli* K-12 W3110.** Active or inactive versions of the CensYBL biosensor containing the wildtype (CBldD) or mutagenized (CBldD*) domain of BldD fused to either N-terminal (NYPet) or C-terminal (CYPet) fragments of the YPet fluorophore were expressed in *E. coli* K-12 W31100 wild type or in strains co-expressing the PDE PdeH or the DGC DgcC from the pCAB18 vector. Cells were grown in LB at 28°C and protein expression was induced for 1 hour using 250 µM IPTG. For each analyzed strain, equal amounts of whole cell proteins were separated using SDS polyacrylamide gel electrophoresis (PAGE). (A) NYPet-CBldD/CbldD* and CYPet-CBldD/CBldD* were detected using polyclonal anti-BldD antibodies (den Hengst *et al.*, 2010) (upper panel) or anti-mCherry antibodies (Invitrogen; lower panel). (B) Loading control for the Western blot shown in panel A. Total proteins were visualized using in-gel 2,2,2-Trichloroethanol (TCE) staining. Following electrophoresis, proteins were cross-linked using UV-light for 5 min and subsequently visualized with transillumination.

**Supplementary Data S1:** Sequence of the c-di-GMP biosensor assembled into pTrc99A-FFA ($Amp^R$)

**key:**

**ypet sequence**
**mCherry sequence**
**sequence coding for the C-terminal domain of BldD**
**linker sequence coding for GSGGG**
***rrnB* terminator**
**restriction sites (HindIII/NdeI)**

>pTrc99A-FFA_*ypet*(split)-*cbldD_mCherry_rrnB* (active CensYBL)

catATGgtctccaagggcgaggagctgttcaccggcgtcgtcccgatcctggtcgagctggacggcgacgtcaacggccacaagt
tctccgtctccggcgagggcgagggcgacgccacctacggcaagctgaccctgaagctgctgtgcaccaccggcaagctgccggtc
ccgtggccgaccctggtcaccaccctgggctacggcgtccagtgcttcgcccgctacccggaccacatgaagcagcacgacttcttc
aagtccgccatgccggagggctacgtccaggagcgcaccatcttcttcaaggacgacggcaactacaagacccgcgccgaggtcaa
gttcgagggcgacaccctggtcaaccgcatcgagctgaagggcatcgacttcaaggaggacggcaacatcctgggccacaagctgg
agtacaactacaactcccacaacgtctacatcaccgccgacaagcagaagaacggcatcaaggccaacttcaagatccgccacaaca
tcgag**ggctccggcggcggc**GAGCCGCCGCCGAAGCTCGTCCTTGACCTGGAGCGCCTCGC
GCACGTCCCCCAGGAGAAGGCGGGCCCGCTCCAGCGCTACGCGGCGACGATCCA
GTCGCAGCGCGGCGACTACAACGGCAAGGTGCTGTCGATCCGTCAGGACGACCT
GCGCACCCTGGCCGTGATCTACGACCAGTCGCCCTCGGTCCTCACCGAGCAGCTG
ATCAGCTGGGGCGTCCTCGACGCGGACGCGCGCCGCGCCGTCGCCCACGAGGAG
AACTGATAAATCGGTCCGGAaggaggATATGAAT**ATGgacaagcagaagaacggcatcaaggcca**
**acttcaagatccgccacaacatcgaggacggcggcgtccagctggccgaccactaccagcagaacaccccgatcggcgacg**
**gcccggtcctgctgccggacaaccactacctgtcctaccagtccaagctgttcaaggacccgaacgagaagcgcgaccacatg**
**gtcctgctggagttcctgaccgccgccggcatcaccgagggcatgaacgagctgtacaagggctccggcggcggc**GAGCC
GCCGCCGAAGCTCGTCCTTGACCTGGAGCGCCTCGCGCACGTCCCCCAGGAGAAG
GCGGGCCCGCTCCAGCGCTACGCGGCGACGATCCAGTCGCAGCGCGGCGACTAC
AACGGCAAGGTGCTGTCGATCCGTCAGGACGACCTGCGCACCCTGGCCGTGATCT
ACGACCAGTCGCCCTCGGTCCTCACCGAGCAGCTGATCAGCTGGGGCGTCCTCGA
CGCGGACGCGCGCCGCGCCGTCGCCCACGAGGAGAACTGATAATGCGACTATCG
**aggagg**TCATTTTTATGgtctccaagggcgaggaggacaacatggccatcatcaaggagttcatgcgcttcaaggtccaca
tggagggctccgtcaacggccacgagttcgagatcgagggcgagggcgagggccgcccgtacgagggcacccagaccgccaag

ctgaaggtcaccaagggcggcccgctgccgttcgcctgggacatcctgtccccgcagttcatgtacggctccaaggcctacgtcaag
cacccggccgacatcccggactacctgaagctgtccttcccggagggcttcaagtgggagcgcgtcatgaacttcgaggacggcgg
cgtcgtcaccgtcacccaggactcctccctgcaggacggcgagttcatctacaaggtcaagctgcgcggcaccaacttcccgtccgac
ggcccggtcatgcagaagaagaccatgggctgggaggcctcctccgagcgcatgtacccggaggacggcgccctgaagggcgag
atcaagcagcgcctgaagctgaaggacggcggccactacgacgccgaggtcaagaccacctacaaggccaagaagccggtccag
ctgccgggcgcctacaacgtcaacatcaagctggacatcacctcccacaacgaggactacaccatcgtcgagcagtacgagcgcgc
cgagggccgccactccaccggcggcatggacgagctgtacaagtgaTAACGGTGGTCCCACCTGACCCCAT
GCCGAACTCAGAAGTGAAACGCCGTAGCGCCGATGGTAGTGTGGGGTCTCCCCA
TGCGAGAGTAGGGAACTGCCAGGCATCAAATAAAACGAAAGGCTCAGTCGAAAG
ACTGGGCCTTTCGTTTTATCTGTTGTTTGTCGGTGAACGCTCTCCTGAGTAGGACA
AATCCGCCGGGAGCGGATTTGAACGTTGCGAACAAACTCCGGGAGGCAGCGTGA
TGCGGCAACAATCACACGGATTTCCCGTGAaagctt

>pTrc99A-FFA_*ypet*(split)-*cbldD**_*mCherry*_*rrnB* (inactive CensYBL; mutations in CbldD are highlighted in red

catATGgtctccaagggcgaggagctgttcaccggcgtcgtcccgatcctggtcgagctggacggcgacgtcaacggccacaagt
tctccgtctccggcgagggcgagggcgacgccacctacggcaagctgaccctgaagctgctgtgcaccaccggcaagctgccggtc
ccgtggccgaccctggtcaccaccctgggctacggcgtccagtgcttcgcccgctacccggaccacatgaagcagcacgacttcttc
aagtccgccatgccggagggctacgtccaggagcgcaccatcttcttcaaggacgacggcaactacaagacccgcgccgaggtcaa
gttcgagggcgacaccctggtcaaccgcatcgagctgaagggcatcgacttcaaggaggacggcaacatcctgggccacaagctgg
agtacaactacaactcccacaacgtctacatcaccgccgacaagcagaagaacggcatcaaggccaacttcaagatccgccacaaca
tcgag**ggctccggcggcggc**GAGCCGCCGCCGAAGCTCGTCCTTGACCTGGAGCGCCTCGC
GCACGTCCCCCAGGAGAAGGCGGGCCCGCTCCAGCGCTACGCGGCGACGATCCA
GTCGCAGGACGGCCGCTACAACGGCAAGGTGCTGTCGATCGACCAGGACCGTCT
GCGCACCCTGGCCGTGATCTACGACCAGTCGCCCTCGGTCCTCACCGAGCAGCTG
ATCAGCTGGGGCGTCCTCGACGCGGACGCGCGCCGCGCCGTCGCCCACGAGGAG
AACTGATAAATCGGTCCGGAaggaggATATGAAT**ATGgacaagcagaagaacggcatcaaggcca**
**acttcaagatccgccacaacatcgaggacggcggcgtccagctggccgaccactaccagcagaacaccccgatcggcgacg**
**gcccggtcctgctgccggacaaccactacctgtcctaccagtccaagctgttcaaggacccgaacgagaagcgcgaccacatg**
**gtcctgctggagttcctgaccgccgccggcatcaccgagggcatgaacgagctgtacaagggctccggcggcggc**GAGCC
GCCGCCGAAGCTCGTCCTTGACCTGGAGCGCCTCGCGCACGTCCCCCAGGAGAAG
GCGGGCCCGCTCCAGCGCTACGCGGCGACGATCCAGTCGCAGGACGGCCGCTAC
AACGGCAAGGTGCTGTCGATCGACCAGGACCGTCTGCGCACCCTGGCCGTGATCT

ACGACCAGTCGCCCTCGGTCCTCACCGAGCAGCTGATCAGCTGGGGCGTCCTCGA
CGCGGACGCGCGCCGCGCCGTCGCCCACGAGGAGAACTGATAATGCGACTATCG
**aggagg**TCATTTTTATGgtctccaagggcgaggaggacaacatggccatcatcaaggagttcatgcgcttcaaggtccaca
tggagggctccgtcaacggccacgagttcgagatcgagggcgagggcgagggccgcccgtacgagggcacccagaccgccaag
ctgaaggtcaccaagggcggcccgctgccgttcgcctgggacatcctgtccccgcagttcatgtacggctccaaggcctacgtcaag
cacccggccgacatcccggactacctgaagctgtccttcccggagggcttcaagtgggagcgcgtcatgaacttcgaggacggcgg
cgtcgtcaccgtcacccaggactcctccctgcaggacggcgagttcatctacaaggtcaagctgcgcggcaccaacttcccgtccgac
ggcccggtcatgcagaagaagaccatgggctgggaggcctcctccgagcgcatgtacccggaggacggcgccctgaagggcgag
atcaagcagcgcctgaagctgaaggacggcggccactacgacgccgaggtcaagaccacctacaaggccaagaagccggtccag
ctgccgggcgcctacaacgtcaacatcaagctggacatcacctcccacaacgaggactacaccatcgtcgagcagtacgagcgcgc
cgagggccgccactccaccggcggcatggacgagctgtacaagtgaTAACGGTGGTCCCACCTGACCCCAT
GCCGAACTCAGAAGTGAAACGCCGTAGCGCCGATGGTAGTGTGGGGTCTCCCCA
TGCGAGAGTAGGGAACTGCCAGGCATCAAATAAAACGAAAGGCTCAGTCGAAAG
ACTGGGCCTTTCGTTTTATCTGTTGTTTGTCGGTGAACGCTCTCCTGAGTAGGACA
AATCCGCCGGGAGCGGATTTGAACGTTGCGAACAAACTCCGGGAGGCAGCGTGA
TGCGGCAACAATCACACGGATTTCCCGTGAaagctt
